# Extension of Colijn-Plazotta tree shape distance metric to unrooted trees

**DOI:** 10.1101/506022

**Authors:** A.A. Morozov

## Abstract

Colijn-Plazotta tree shape labeling scheme allows to describe an arbitrary phylogenetic tree topology by recursively labeling all nodes from tips to root with integers. The multisets of these labels can then be used to estimate the difference between topologies using eg Euclidean distance. In this work I propose an extension of the labeling scheme (and thus a distance metric) to unrooted trees, which is achieved by labeling all rooted subtrees within a given tree. To avoid exhaustively enumerating the subtrees, the labels are collected into a dependency graph and calculated in a single pass. A proof-of-concept implementation is available at https://github.com/synedraacus/metrics.

---

Analysis of the phylogenetic tree shapes, which ignores the leaf labels and concentrates on the topology itself, can be used to study the large-scale evolutionary patterns [Moers, Heard 1997]. Usually such works center on estimating one or several shape parameters, most commonly balance, on a set of trees [Purvis et al., 2011; Blum, Francois 2006]. Although useful for estimating various parameters of evolution and applicable to the large-scale data, variation in tree balance or any other parameter is by definition one-dimensional: it can only tell a researcher that some subset of trees is more balanced than another. More complex analyses such as clustering trees into an unknown number of differently-evolving groups, exploratory analysis via *eg* MDS or detecting *a priori* unknown patterns of shape variation require a more complex mathematical apparatus. A proper distance metric between two arbitrary trees which is symmetric, equals zero only for equal trees, and satisfies the triangle inequality was proposed in a recent paper [Colijn, Plazotta 2018].

It is based on labeling all the nodes in the tree with positive integers in the following way: first, leaf nodes (or, equivalently, single-leaf trees) are labeled 1. Then, each internal node is treated as *(k, j)*-node, where *k* and *j* are labels of its children. The label of a (*k, j*)-subtree is equal to a position of this tuple in a lexicographically sorted list of all integer pairs, *ie* (1), (1, 1), (2, 1), (2, 2), (3, 1), (3, 2)… and so on. Since (*k, j*) and (*j, k*)-nodes are equivalent from a phylogenetic point of view, a convention is to assume that *k>j*. For example, an (1, 1)-node, a parent of two leaves, is given label 2 because it is second in this list; a (2, 1) node, a parent of (1, 1) and another leaf, is given label 3 and so on. A label for arbitrary *k* and *j* is given by the following formula:

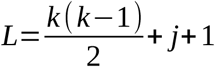

With this formula, a simple walk from leaves to root can label all the nodes in a tree, and the root label is sufficient to uniquely describe the tree shape. However, comparing root labels is not useful from a biological point of view. For example, an addition of a single leaf will lead to a significant increase of root label, although the rest of its topology remains exactly the same. Instead, the authors propose describing the tree with a complete list of its labels, which is redundant, but includes the information about all subtrees present in a tree. This list is converted into a multiset, where each label *L* is associated with its occurrence count *k*L, and the Euclidean distance is calculated between these counts in two trees. Although occurrence counts are not, strictly speaking, independent (each label above 1 implies a certain multiset of labels among its descendants), Euclidean distance is a proper distance metric and was shown to work well in practice. Authors have successfully used it to distinguish both between the viral trees from different environments and between simulated trees produced by different algorithms [Colijn, Plazotta 2018].

One disadvantage of this labeling scheme is that it assumes a rooted tree. Real trees, on the other hand, usually are inherently unrooted because most common substitution models are time-reversible. While a plausible root can be found using outgroups, molecular clock assumption, nonreversible substitution models [Huelsenbeck, Bollback, Levine 2002], midpoint rooting [Hess, De Moraes Russo 2007] and other approaches, these methods are not always reliable or even available. Re-rooting or arbitrarily rooting the tree can in some cases be irrelevant from the biological point of view, but it does affect the label multiset. Thus, a variant of Colijn-Plazotta labeling applicable to unrooted trees can be useful in practice.

Such a variant, based on labeling all rooted subtrees in a given unrooted tree, is described below. It’s important to note that this approach considers rooted *sub*trees that can be produced by splitting the original tree across one of the edges, not possible rootings of the entire tree (which would introduce additional nodes and thus additional labels). The naïve solution would be to produce these subtrees and calculate their labels. But the subtrees are largely overlapping, which makes enumerating them both computationally inefficient, since every value is included in multiple subtrees, and mathematically nontrivial, requiring the procedure for correctly joining label multisets. Instead, all values can be mapped on an original unrooted tree using the following procedure.

Any node in an unrooted binary tree *T* has either one or three edges, depending on whether it’s a leaf or an internal node. Subtrees that are connected to the node in question by these edges are themselves binary trees, and therefore can be considered rooted at this node and described by a single root label each. These labels are, in turn, produced by recursively annotating the node’s neighbours, neighbours of their neighbours, and so on until the leaves. In this way every node in the tree can be annotated with a vector of either one or three labels (Fig. 1a).

**Fig. 1:**
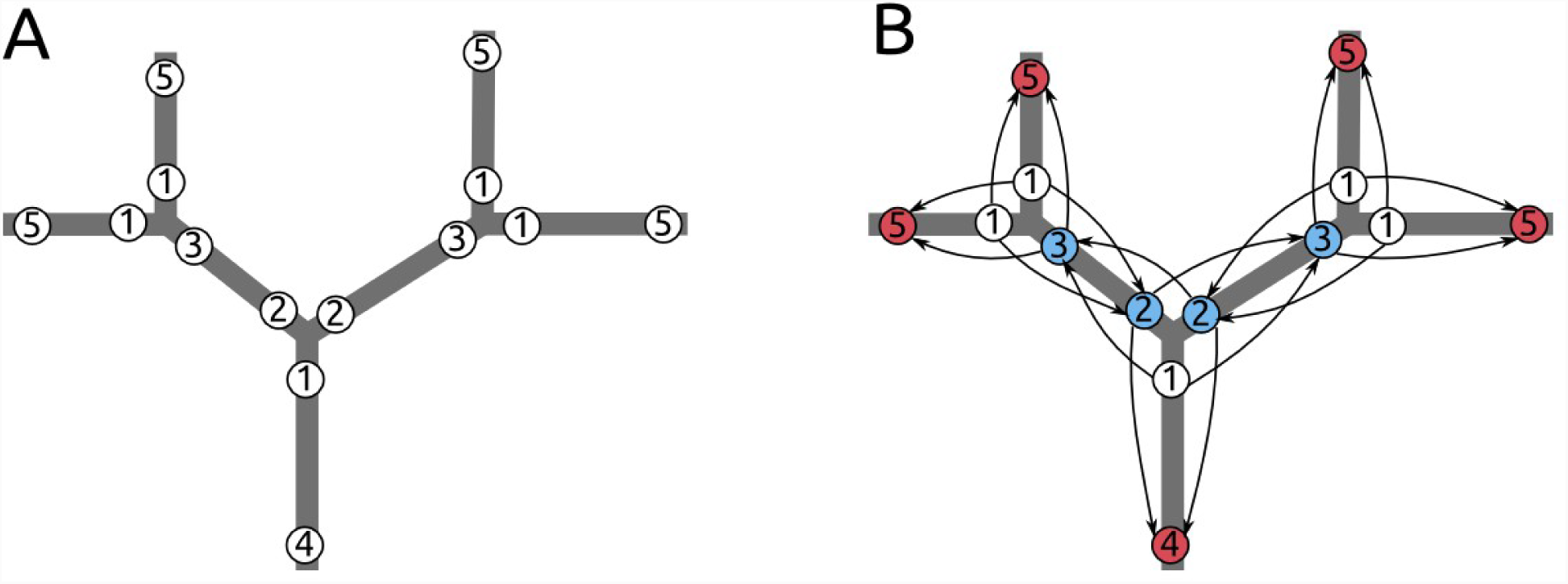
An example of unrooted tree labeling. (A): The labels on a five-leaf tree. (B): The label graph on the same tree. Starting nodes (branches leading to leaves) are colored white, intermediate nodes are blue, and terminal nodes with zero outdegree are red.

If every node in a tree is annotated in this way, the multiset of their labels *L* includes a label multiset of every subtree. This can be proven by interpreting values as attached to the edges, rather than nodes. The equivalence of these interpretations can, in turn, be shown by a graph *M* whose nodes are corresponding to nodes and edges of *T*, and edges connect the nodes whose corresponding elements are incident in *T*. The labels are not just attached to the nodes of *T*: each of them by definition describes the subtree to which this node is connected by exactly one of the edges. Since the label corresponds to exactly one node and exactly one incident edge in *T*, it corresponds to a unique edge in *M*. If the labeling is complete, *ie* every node in *T* has every label that it can have, then all edges in *M* have a value. It is then trivial, for a given *M*, to construct *T’* which has the same topology as *T*, but each edge (instead of node) has a pair of labels. The label multisets of *T* and *T’* are identical: the values are only moved from nodes to edges or *vice versa*, but not created or lost. Since *T* and *T’* can be converted to each other via *M*, they are equivalent for the purposes of this work.

Consider any two rooted subtrees of *T’*, produced by splitting it across one of the edges. Their root labels are labels on this edge and therefore are included in the label multiset of *T’*. Each label within a subtree is laying on one of its edges, which by definition are also edges of *T’*, and can be similarly shown to belong to the label multiset of *T’* and therefore the label multiset of *T*.

To actually produce this multiset, the labels are connected into a directed graph where the nodes are labels and incoming edges at each node connect it with nodes that are necessary to calculate a label (fig. 2b). In other word, they correspond to the branches of one of the subtrees of *T* directed leaf-to-root. Leaf labels (white on figure) are set to one by definition of the CP labeling, and the graph is traversed from them to produce the rest of the labels.

Proof-of-concept Python3.6 implementation (using gmpy2, a GMP wrapper, for handling large numbers) is available at https://github.com/synedraacus/metrics. It should be noted that extending the method to unrooted trees has lead to a significant increase in RAM cost. The original rooted version already has high requirements due to the quick increase of label values, but they can be managed by hashing nodes’ labels the moment their parents’ labels are set, and analyzing the frequencies of label hashes instead of the label frequencies. In this way, only a small number of labels has to be stored at any given moment, which decreases memory consumption drastically at the cost of risking hash collision (which is usually negligible). This method is less effective for unrooted trees: since the label graph is bigger than an underlying tree, as well as more densely connected, more labels need to be kept unhashed simultaneously. In practice, processing a single tree with 400 leaves requires approx. 50 Gb of RAM.

## Acknowledgements

The reported study was funded by RFBR according to the research project ? 18-34-00441.

## Literature

Blum M.G.B., Francois O. Which random processes describe the tree of life? A large-scale study of phylogenetic tree imbalance. Systematic Biology, 2006, 55(4): 685–691.

Colijn C., Plazotta G. A metric on phylogenetic tree shapes. Systematic Biology, 2018, 67(1): 113–126.

Hess P.H., De Moraes Russo C.A. An empirical test of the midpoint rooting method. Biological Journal of the Linnean society, 2007, 92, 669–674.

Huelsenbeck J.P., Bollback J.P., Levine A.M. Inferring the root of a phylogenetic tree. Systematic Biology, 2002, 51(1): 32–43.

Mooers A.Ø., Heard S.B. Inferring evolutionary processes from phylogenetic tree shape. The Quarterly Review of Biology, 1997, 72(1): 31–54.

Purvis A., Fritz S.A., Rodríguez J., Harvey P.H., Grenyer R. The shape of mammalian phylogeny: patterns, processes and scales. Philos Trans R Soc Lond B Biol Sci, 2011, 366(1577): 2462–2477.

